# Computational and experimental analyses of alanine racemase suggest new avenues for developing allosteric small-molecule antibiotics

**DOI:** 10.1101/2022.10.29.514358

**Authors:** Arie Van Wieren, Jacob D Durant, Sudipta Majumdar

**Affiliations:** Madia Department of Chemistry, Biochemistry, Physics and Engineering, Indiana University of Pennsylvania, Indiana, PA 15705; Department of Biological Sciences, University of Pittsburgh, Pittsburgh, PA 15260; The Johns Hopkins University School of Medicine, Baltimore, MD 21205

**Author notes:** Correspondence: Sudipta Majumdar, Madia Department of Chemistry, Biochemistry, Physics and Engineering, Indiana University of Pennsylvania, Indiana, PA 15705. Correspondence: Jacob Durrant, Department of Biological Sciences, University of Pittsburgh, Pittsburgh, PA 15260.

**Keywords:** Alanine racemase, antibacterial resistance, drug discovery, protein engineering, enzyme kinetics, protein structure-function, computer-aided drug design, computational biology

## Abstract

Given the ever-present threat of antibacterial resistance, there is an urgent need to identify new antibacterial drugs and targets. One such target is alanine racemase (Alr), an enzyme required for bacterial cell-wall biosynthesis. Alr is an attractive drug target because it is essential for bacterial survival but is absent in humans. Here, we investigate the Alr from *M. tuberculosis* (MT), the pathogen responsible for human tuberculosis, as a model Alr enzyme. MT-Alr functions exclusively as an obligate homodimer formed by two identical monomers. Both monomers contribute to the overall composition of their active sites. Therefore, disrupting the dimer interface could inhibit MT-Alr activity. Using computational methods, we identified seven interfacial residues predicted to be responsible for MT-Alr dimerization. Mutating one of the seven residues, Lys261, to alanine resulted in a completely inactive enzyme. Further investigation suggested a potential drug-binding site near Lys261 that might be useful for allosteric drug discovery.

**Summary:** The bacterial protein alanine racemase (Alr) converts L-alanine to D-alanine, a critical component of the bacterial cell wall. Cycloserine, a known antibiotic, inhibits Alr by binding to the same pocket that alanine binds. Several human proteins have similar pockets, so cycloserine has severe side effects. We identified additional Alr pockets and discovered that altering one of them abolishes Alr activity. Molecules that bind this pocket may similarly impact Alr activity, helping to address the ongoing antibiotic resistance crisis.

## Introduction

Antibacterial resistance is among the most serious challenges to global public health and food production. Addressing this threat requires preventing infections, restricting the development of resistance through prudent use of existing antibiotics, halting the spread of resistance when it does develop, and developing novel pharmaceuticals that are not yet subject to resistance. Alarmingly, no new antibacterial class has been introduced since 2000, when the oxazolidinone linezolid was approved.^1^ Strategies for overcoming this discovery gap include repurposing old classes of antibiotics/adjuvants, turning to antibiotic combination therapy,^2-4^ and developing new antibiotics that act through novel/underutilized drug targets.

Alanine racemase (Alr), a PLP-dependent enzyme that plays an essential role in cell wall synthesis, is one potential drug target. Alr racemizes L-alanine to D-alanine, a key building block in peptidoglycan biosynthesis.^5,6^ It is an attractive antibacterial target because it is essential for bacterial survival but is absent in humans. Prior research identified structural analogs of D-alanine as Alr inhibitors, leading to the discovery of the FDA-approved drug cycloserine (brand name Seromycin).^7-9^ Unfortunately, like other inhibitors that target the Alr active site, cycloserine lacks specificity because it acts on other PLP-containing enzymes in humans, especially in the central nervous system (CNS).^10,11^ Given this CNS activity, recent studies have suggested using cycloserine to treat psychiatric diseases such as schizophrenia, anxiety disorder, major depression, autism, and dementia.^12,13^ However, severe side effects (e.g., confusion, seizure, speech disorder, dizziness, coma) limit its utility as an antibiotic drug. These challenges suggest the need for allosteric Alr inhibitors that do not target the PLP-dependent active site, enabling safer inhibition.

Alr crystal structures from different organisms have been recently resolved, revealing much about this enzyme’s tertiary structure and catalytic activity. With few exceptions, Alrs are homodimers formed by the head-to-tail association of two monomers.^14-17^ Both monomers are essential because they contribute to the overall composition of the active site, the substrate entryway, and the binding pocket.^18^ In theory, molecules that disrupt the dimer interface could inhibit Alr activity, a strategy that has been successfully applied to other drug targets such as HIV protease, caspase, etc.^19-22^ Many of the residues that participate in the dimer interface are also highly conserved, suggesting the possibility of broad-spectrum dimer-disrupting antibiotics.^18^

In this work, we use mutagenesis to identify an Alr salt bridge distant from the active site that may be critical for dimer association. Mutating one of the salt-bridge residues to alanine abolishes catalytic activity, presumably because of impaired dimerization. We also use computational methods to show that the salt bridge is adjacent to a previously unknown druggable pocket, suggesting a pharmaceutical strategy for disrupting dimerization by interfering with a critical anchor point.

### Materials and Methods

All reagents and chemicals used as buffers and substrates of the highest grade chemically available were purchased from Sigma-Aldrich, Fisher, Acros, or Alfa Aesar. UV-Vis spectrophotometric data were collected using Cary 60 UV-Vis Spectrophotometer (Agilent) and Multiskan GO (Fisher Scientific). Protein purifications were conducted using BioLogic DuoFlow (Bio-Rad). Alr from *M. tuberculosis* (MT-Alr) and L-alanine dehydrogenase from *S. coelicolor* (Ald) were cloned, expressed, purified, and characterized in the lab.^23,24^

#### Predicting residues critical for dimerization

We followed a multi-step protocol to identify candidate interfacial residues critical for dimerization. First, we used visual molecular dynamics (VMD)^25^ to identify Alr residues that came within 4 Å of the opposite monomer but were at least 8 Å from the catalytic site. The 6SCZ^26^ crystal structure was used for this analysis because its active site is fully occupied (i.e., PLP is conjugated to cycloserine), and we used this conjugate to judge distances to the catalytic site. Second, we used the Consurf server^27^ to identify those residues with above-average conservation scores. Third, we applied Robetta,^28^ a program for computational alanine scanning, to the 6SCZ structure and retained residues predicted to decrease the binding energy by at least 1.0 kcal/mol when mutated to alanine. Finally, we used BINANA^29^ and visual inspection to identify residues that participated in specific (e.g., salt-bridge, hydrogen-bond, π-stacking) interactions with opposite-monomer residue(s).

#### Mutagenesis

The plasmid pET28b-MT_Alr^24^ served as the DNA template to generate seven MT-Alr single mutants (Glu72Ala, Asp135Ala, Asn139Ala, Lys261Ala, Glu267Ala, Arg371Ala, and Arg373Ala) using a Q5 site-directed mutagenesis kit (NEB). The corresponding primers were designed using the NEBase Changer tool (http://nebasechanger.neb.com/; Table 1). The PCR mixture (10 μL) contained 1 μL of 10 ng/μL template DNA plasmid, 1 μL each of 5 μ M forward and reverse primers, 5 μL of Q5 HS master mix, and 2 μL of ddH_2_O. The PCRs were performed in a BioRad T100 Thermal Cycler with the following parameters: 98 °C for 30 s followed by 25 cycles of 98 °C for 10 s, T_a_ °C (suggested annealing temperature for the primers) for 20 s, and 72 °C for 3 m, and a final extension time of 2 m at 72 °C. The PCR reactions were confirmed by DNA gel electrophoresis. The KLD reactions were performed at room temperature for 10 min with 0.5 μL of amplified PCR product, 2.5 μL of KLD reaction buffer, 1.5 μL of ddH_2_O, and 0.5 μL of KLD enzyme mixture. 2.5 μL of KLD mixtures were chemically transformed into 5-alpha competent *E. coli* cells. The mutations were confirmed by DNA sequencing (Genewiz).

**Table 1.**
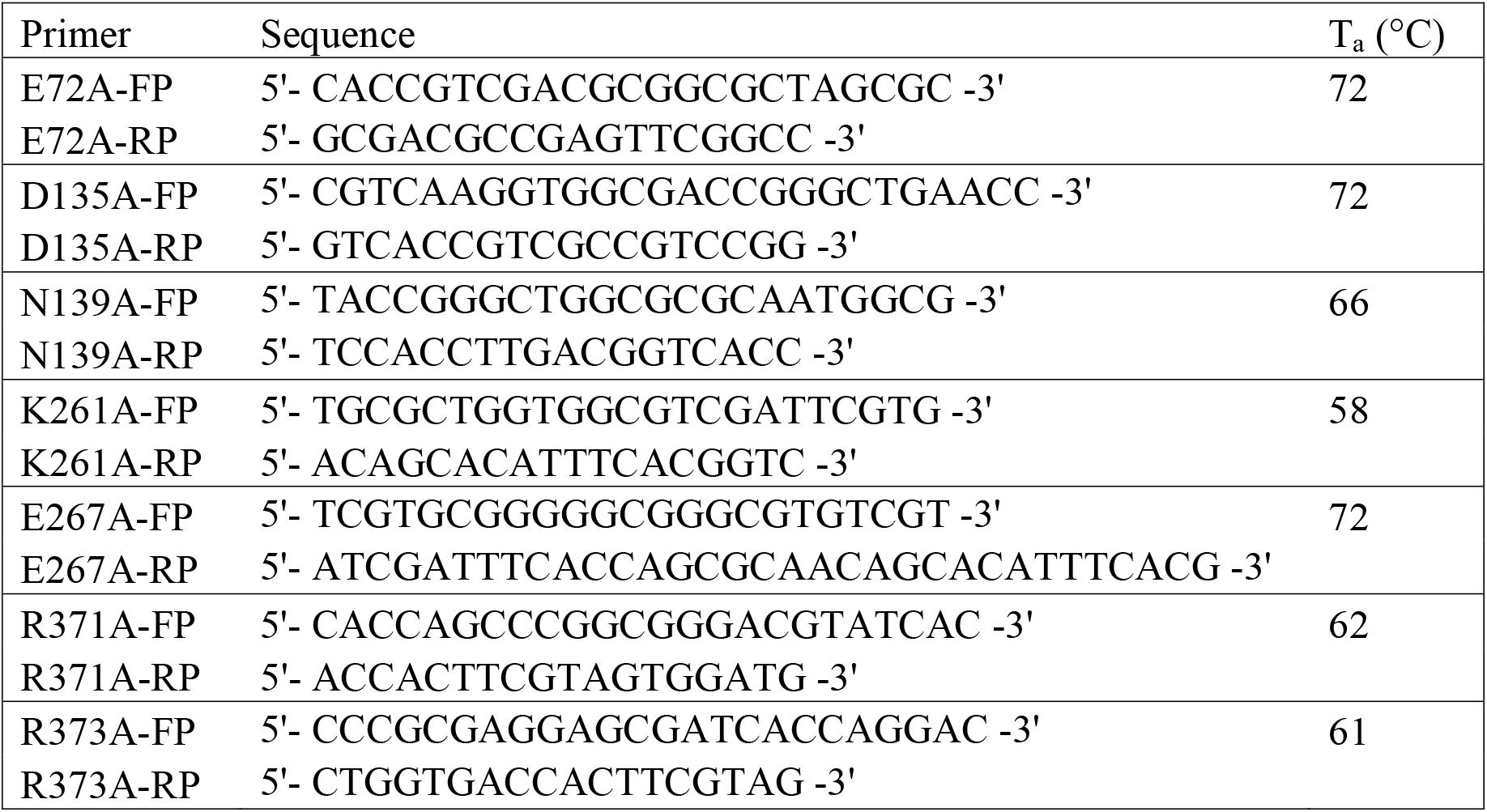
Primers used for single-alanine mutant generation with corresponding annealing temperatures (T_a_).

#### Protein expression and purification

The MT-Alr mutant proteins were cloned with an N-terminal His-tag and were expressed in *E. Coli* BL21 (DE3) cells. A typical large-scale purification involved a bacterial culture grown in 2 × 1 L of terrific broth (TB) medium with 50 μ /mL kanamycin. This broth was inoculated with 5 mL starter culture and shaken at 180 rpm at 37 °C until the OD_600_ reached 0.5. The incubator temperature was then decreased to 30 °C and shaken until the OD_600_ reached 0.6, at which point expression was induced to the final concentration of 1 mM isopropyl-β-D-thiogalactoside (IPTG). The culture was incubated for a further 24 hours at 30 °C with shaking at 180 rpm. Cells were harvested by centrifugation at 7,000 rpm for 5 min. The cell pellets were dissolved in 20 mL of cold buffer (20 mM Tris, 100 mM NaCl, pH 8.0) with 20 μL Halt Protease Inhibitor Cocktail (100x). Cells were lysed by sonication (10 min, 5 s on and 5 s off), and the lysate was cleared by centrifugation. The supernatant was applied to a 1 mL IMAC column (Bio-Rad) and eluted with a linear gradient (100 mL) of 250 mM imidazole buffered with 20 mM Tris and 100 mM NaCl (pH 8.0). Fractions containing pure protein were collected and dialyzed against 20 mM Tris-HCl, 100 mM NaCl (pH 8.0), and then stored at −80 °C. The purity of the protein fractions was verified by SDS-PAGE (Fig S1, supporting information).

#### Enzyme characterization

Following the optimal enzymatic assay condition determined for the wild-type MT-Alr enzyme,^24^ the mutant enzyme activities were measured in the D-to L-alanine direction. We monitored the production of NADH (ε_340_ = 6.22 mM^−1^ cm^−1^) at 340 nm (30 °C, pH 9.0) as the L-alanine was converted to pyruvate and ammonia by L-alanine dehydrogenase (Scheme S1, supporting information). The reaction mixture (250 μL) contained 0.01 − 50 mM D-alanine (dissolved in buffer), NAD (906 μM), L-alanine dehydrogenase (2.76 μ M), and mutant MT-Alr (0.6 μ M). The steady-state kinetic constants were determined by fitting the kinetic data (triplicate) to the Michaelis-Menten equation using GraphPad Prism7 (GraphPad Software, Inc.).

#### Homology modeling and structural alignment

Homology models of Alrs from *Enterococcus faecium* (EF-Alr), *Neisseria gonorrhoeae* (NG-Alr), and *Klebsiella pneumoniae* (KP-Alr) were generated using the target-template sequence-alignment tool available through the SWISS-MODEL Workspace (https://swissmodel.expasy.org/; Fig S2, supporting information). We used Chimera MatchMaker^30^ to structurally align these three models as well as two crystal structures of Alrs from *M. tuberculosis* (MT-Alr; 1XFC^31^) and *Streptococcus pneumoniae* (SP-Alr, 3S46^18^).

#### Predicting druggable pockets

To predict druggable pockets, we first processed a crystal structure of MT-Alr (PDB code: 1XFC)^31^ using Maestro 13.2.128 (Schrodinger 2022-2). Missing internal and terminal residues in each of the monomers were constructed using Prime,^32,33^ and the newly built protein structure was preprocessed using the Protein Preparation Wizard.^34^ We then removed all water molecules from the energy-minimized structure and analyzed it for potential druggable pockets using Schrödinger’s SiteMap program^35^; the FTMap web server^36^; and FPocketWeb,^37^ a web-based implementation of fpocket.^38,39^

## Results and Discussion

The alanine racemase from *M. tuberculosis* (MT-Alr) is an ideal model for the present study because much prior research has focused on its structure, both with and without inhibitors bound to the active site.^26,31^ Like most other Alr enzymes, MT-Alr is a functionally obligate homodimer with a conserved dimerization motif.^24,31^ Disruption of this motif should therefore inhibit its activity.

### Identifying residues critical for dimerization

Protein-protein interfaces are important but challenging targets for inhibitor design. A crucial first step is identifying potential sites along these interfaces where interface-perturbing drugs might bind and disrupt critical interactions. These sites are typically associated with the small subset of all interfacial residues that contribute substantially to the overall protein binding energy.^40^

Using computational and structural analyses, we identified eight candidate interfacial residues likely to contribute to dimerization: Arg140, Tyr271, Asp320, Asn139, Lys261, Arg371, Glu365, and Arg373. We selected these residues based on four criteria. (1) The residues line the dimeric interface but are distant from the catalytic pocket and so are unlikely to impact catalysis directly. (2) Each residue participates in specific interactions with residue(s) of the opposite monomer (e.g., salt bridges and hydrogen bonds, not just hydrophobic contacts). (3) The residues are at least modestly conserved among Alr proteins across many species, suggesting they play an essential role in protein function. (4) Computational alanine scanning suggests each residue contributes at least 1.0 kcal/mol to the binding energy.

Of these eight residues, we selected four that were distributed roughly evenly across the dimeric interface: Asn139, Lys261, Arg371, and Arg373. To this list of four, we added Asp135, which forms a salt bridge with Lys261; Glu72, which forms a salt bridge with Arg373; and Glu267, which forms a hydrogen bond with Asn139. These three additional residues also line the dimeric interface, are distant from the catalytic site, and are at least modestly conserved.

The seven selected residues can be divided into three clusters based on their locations and interaction patterns. Two clusters, 1A and 1B, are identical and located at the front and back ends of the dimer interface. Cluster 2 is in the middle of the interface (Fig 1 and Table 2).

**Table 2.**
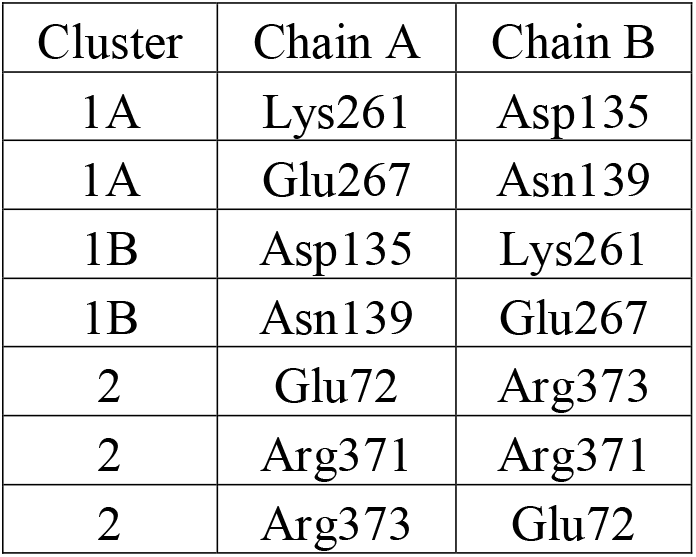
Potential dimer-promoting clusters at the MT-Alr dimer interface.

| Cluster | Chain A | Chain B |
| --- | --- | --- |
| 1A | Lys261 | Asp135 |
| 1A | Glu267 | Asn139 |
| 1B | Asp135 | Lys261 |
| 1B | Asn139 | Glu267 |
| 2 | Glu72 | Arg373 |
| 2 | Arg371 | Arg371 |
| 2 | Arg373 | Glu72 |

**Fig 1.**
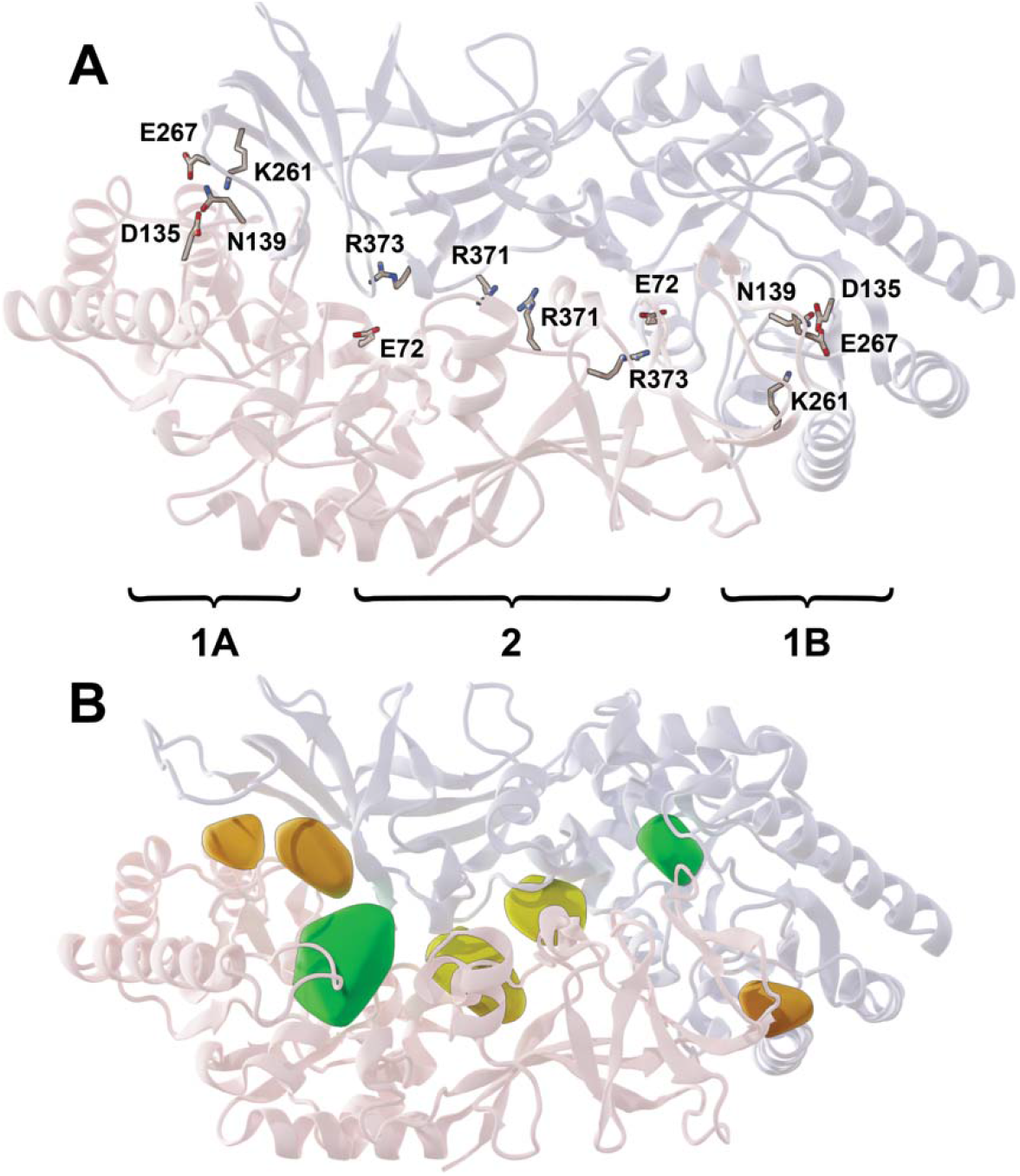
MT-Alr homology model. (A) The two monomers of the Alr dimer are shown in pink and blue ribbon, respectively. Predicted critical residues are shown in stick representation with text labels. The three hotspot clusters are labeled with braces. (B) FTMap-identified druggable hotspots near the Asp135/Lys261 salt bridge (orange), orthosteric cycloserine-binding site (green), and central Arg371/Arg371 juxtaposition (yellow).

### Verifying predicted dimer-promoting residues using mutagenesis and enzymatic assays

We used mutagenesis to evaluate the impact of Glu72, Asp135, Asn139, Lys261, Glu267, Arg371, and Arg373 on catalytic activity. Because Alr is an obligate homodimer, each residue is present on both sides of the interface, so each mutation had a twofold impact. We generated a single-point alanine mutant for each of the seven potential dimer-promoting residues. We chose alanine because it eliminates the amino acid side chain beyond the β carbon but does not alter the main-chain conformation (as can glycine and proline).

We determined the steady-state rate constants for all mutant proteins under the same conditions used for wild-type MT-Alr^24^ (Table 3). To examine the impact of the Glu72/Arg373 salt bridge, we considered the Glu72Ala and Arg373Ala mutants. Both exhibited similar k_cat_ and K_m_ values as the wild-type, suggesting the Glu72/Arg373 salt bridge is not critical for catalytic activity or dimerization.

**Table 3.**
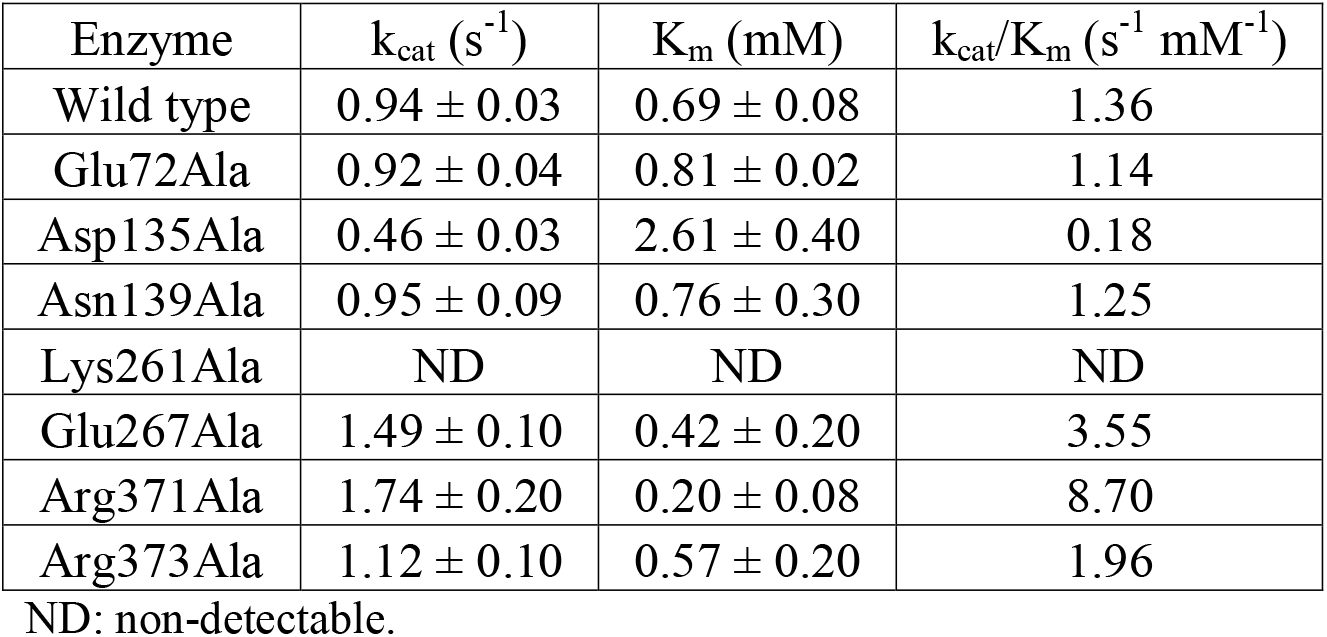
Enzymatic rate constants for MT-Alr wild-type and single-alanine mutants.

| Enzyme | $k_{\text{cat}}$ ($\text{s}^{-1}$) | $K_{\text{m}}$ (mM) | $k_{\text{cat}}/K_{\text{m}}$ ($\text{s}^{-1} \text{mM}^{-1}$) |
| --- | --- | --- | --- |
| Wild type | $0.94 \pm 0.03$ | $0.69 \pm 0.08$ | 1.36 |
| Glu72Ala | $0.92 \pm 0.04$ | $0.81 \pm 0.02$ | 1.14 |
| Asp135Ala | $0.46 \pm 0.03$ | $2.61 \pm 0.40$ | 0.18 |
| Asn139Ala | $0.95 \pm 0.09$ | $0.76 \pm 0.30$ | 1.25 |
| Lys261Ala | ND | ND | ND |
| Glu267Ala | $1.49 \pm 0.10$ | $0.42 \pm 0.20$ | 3.55 |
| Arg371Ala | $1.74 \pm 0.20$ | $0.20 \pm 0.08$ | 8.70 |
| Arg373Ala | $1.12 \pm 0.10$ | $0.57 \pm 0.20$ | 1.96 |

ND: non-detectable.

To examine the impact of the Asn139/Glu267 interaction, we considered the Asn139Ala and Glu267Ala mutants. Here the effect was less straightforward. Paradoxically, the Asn139Ala mutation had minimal impact on catalytic activity, but the Glu267Ala mutation increased k_cat_ and decreased K_m_ (i.e., improved catalytic efficiency).

We also used mutagenesis to explore the impact of the two juxtaposed Arg371/Arg371 residues at the center of the Alr dimer. The Arg371Ala mutant resulted in a six-fold increase in catalytic efficiency. It is surprising that two positively charged amino acids are so juxtaposed. We hypothesize that these two residues might reciprocally regulate enzyme catalysis. *M. tuberculosis* requires only small amounts of D-alanine for survival (5-10 μg/mL)^41^; excessive conversion of L-alanine to D-alanine could in theory deplete the L-alanine pool available for protein synthesis and other critical cellular functions. Perhaps MT-Alr is too catalytically active unless the Arg371/Arg371 juxtaposition is present to reduce—but not eliminate—Alr’s propensity for dimerization. Alternatively, perhaps the two Arg371/Arg371 residues, together with the conserved residues Glu365/Glu365, form an allosteric site (Fig 2) that allows an endogenous ligand to reciprocally regulate Alr catalysis. If such a ligand were to weaken the interaction between Glu365 and Arg371, it might reduce catalytic activity by discouraging dimerization just as the Arg371Ala mutation apparently does. While we are aware of no such endogenous allosteric effector, others have found evidence of Alr allostery^42^ and have even proposed allosteric sites near the Arg371/ Arg371 juxtoposition.^43^

**Fig 2.**
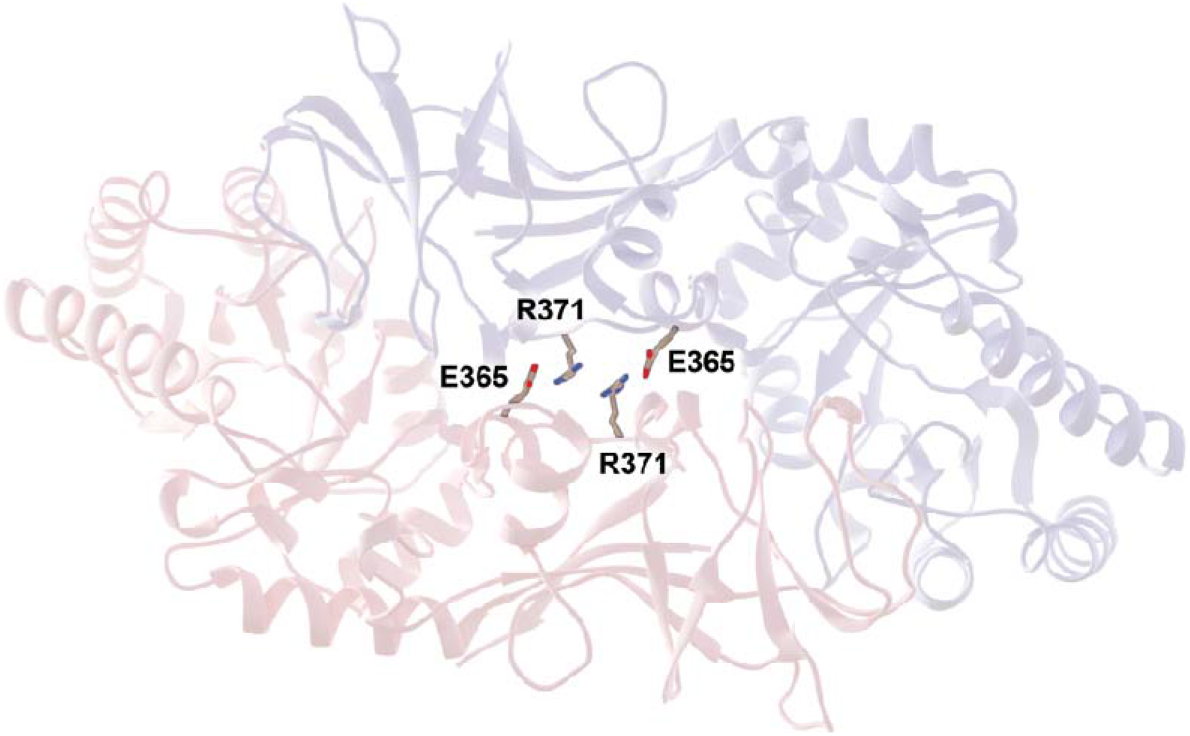
Locations of key residues near the center of the MT-Alr dimer (PDB: 1XFC). These residues participate in electrostatic interactions that may impact enzyme catalysis.

Finally, to examine the impact of the Asp135/Lys261 salt bridge, we considered the Asp135Ala and Lys261Ala mutants. The Asp135Ala mutation reduced k_cat_ and increased K_m_, resulting in an 8-fold decrease in catalytic activity. The Lys261Ala mutation resulted in a complete loss of catalytic activity. Given that these residues are distant from the catalytic pocket, we hypothesize that the Asp135/Lys261 salt bridge is critical for dimerization. Its disruption may indirectly impede catalysis by reducing MT-Alr dimerization.

### Identification of druggable targets

To assess whether carefully designed small-molecule ligands could mimic the effect of these mutations on Alr dimerization, we used three methods to search for druggable Alr pockets. First, we used the FTMap server^36^ to computationally dock a set of small molecular fragments with drug-like chemical properties onto the entire surface of the MT-Alr (1XFC)^31^ structure. Locations on the surface where fragments tend to congregate are known as druggable hotspots and often correspond to known ligand-binding pockets. FTMap identified eleven hotspots that can be generally divided into three groups. Five of the eleven hotspots fell near one of the two known cycloserine-binding catalytic sites, providing a positive control (Fig 1B, in green). FTMap automatically removes non-standard residues, so this pocket did not include the conjugated PLP molecule of the original structure. Three additional hotspots were located near the identified Asp135/Lys261 salt bridge (Fig 1B, in orange). Compounds that bind at this position could potentially disrupt the salt bridge, discouraging dimerization and, thus, catalysis. Finally, three hotspots were located near the central Arg371/Arg371 juxtaposition (Fig 1B, in yellow). Given that mutating these residues to alanine improves catalytic activity (presumably by encouraging dimerization), compounds that shift these residues into a conformation even less amenable to dimerization could reduce Alr activity.

As a second assessment of Alr druggability, we used Schrödinger’s SiteMap program^35^ to identify druggable locations on the MT-Alr (1XFC)^31^ protein surface. SiteMap identified five potential pockets. Of these, one corresponded to a cycloserine-binding catalytic site (Fig S3, in green), even though SiteMap retained the conjugated PLP bound in that site. This finding again serves as a positive control. Three of the five predicted sites were again adjacent to the Asp135/Lys261 salt bridge (Fig S3, in orange), including the top-ranked pocket, and one of the five corresponded to the Arg371/Arg371 juxtaposition (Fig S3, in yellow).

As a third assessment of Alr druggability, we used FPocketWeb,^37^ a web-based implementation of fpocket,^38,39^ to identify MT-Alr (1XFC)^31^ cavities. Among the top eight FPocketWeb-identified pockets, the top two corresponded to a cycloserine-binding catalytic site (Fig S4, in green), which did not include a bound PLP due to FPocketWeb processing. Two predicted sites were again adjacent to the Asp135/Lys261 salt bridge (Fig S4, in orange), and one corresponded to the Arg371/Arg371 juxtaposition (Fig S4, in yellow).

### An extended-spectrum antibiotic drug target

To assess whether dimer-disrupting compounds that bind near the Asp135/Lys261 salt bridge might serve as extended-spectrum antibiotics, we used ClustalW to align the amino acid sequences of the five Alrs from different pathogenic bacterium listed as the highest antibiotic resistance threats in the United States by the Centers for Disease Control and Prevention. The pathogens include drug-resistant *M. tuberculosis* (MT-Alr), gram-positive Vancomycin-resistant *Enterococcus faecium* (EF-Alr), gram-positive multidrug-resistant *Streptococcus pneumoniae* (SP-Alr), gram-negative multidrug-resistant *Neisseria gonorrhoeae* (NG-Alr), and gram-negative Carbapenem-resistant *Klebsiella pneumoniae* (KP-Alr).

As a control, we first considered the residues involved in catalytic alanine racemization (boxed in blue). These are conserved in all Alrs, as expected given their central importance to the catalytic mechanism. Next, we considered the residues analogous to Asp135/Lys261 (boxed in red). These are either completely (Asp) or semi-completely (Lys/Arg) conserved in all Alrs (Fig 3). Structural alignment of these Alrs further supports spatial conservation (Fig 3). This conservation suggests that the Asp135/Lys261 salt bridge is not unique to MT-Alr; a similar salt bridge may be critical for dimerization and catalytic efficiency in other species as well. Compounds that disrupt Alr dimerization in *M. tuberculosis* may therefore be effective against other bacterial Alrs, including those from gram-positive, gram-negative, and mycobacteria. If this potential mechanism of action is ubiquitous, it could enable the discovery of novel broad-spectrum antibiotics.

**Fig 3.**
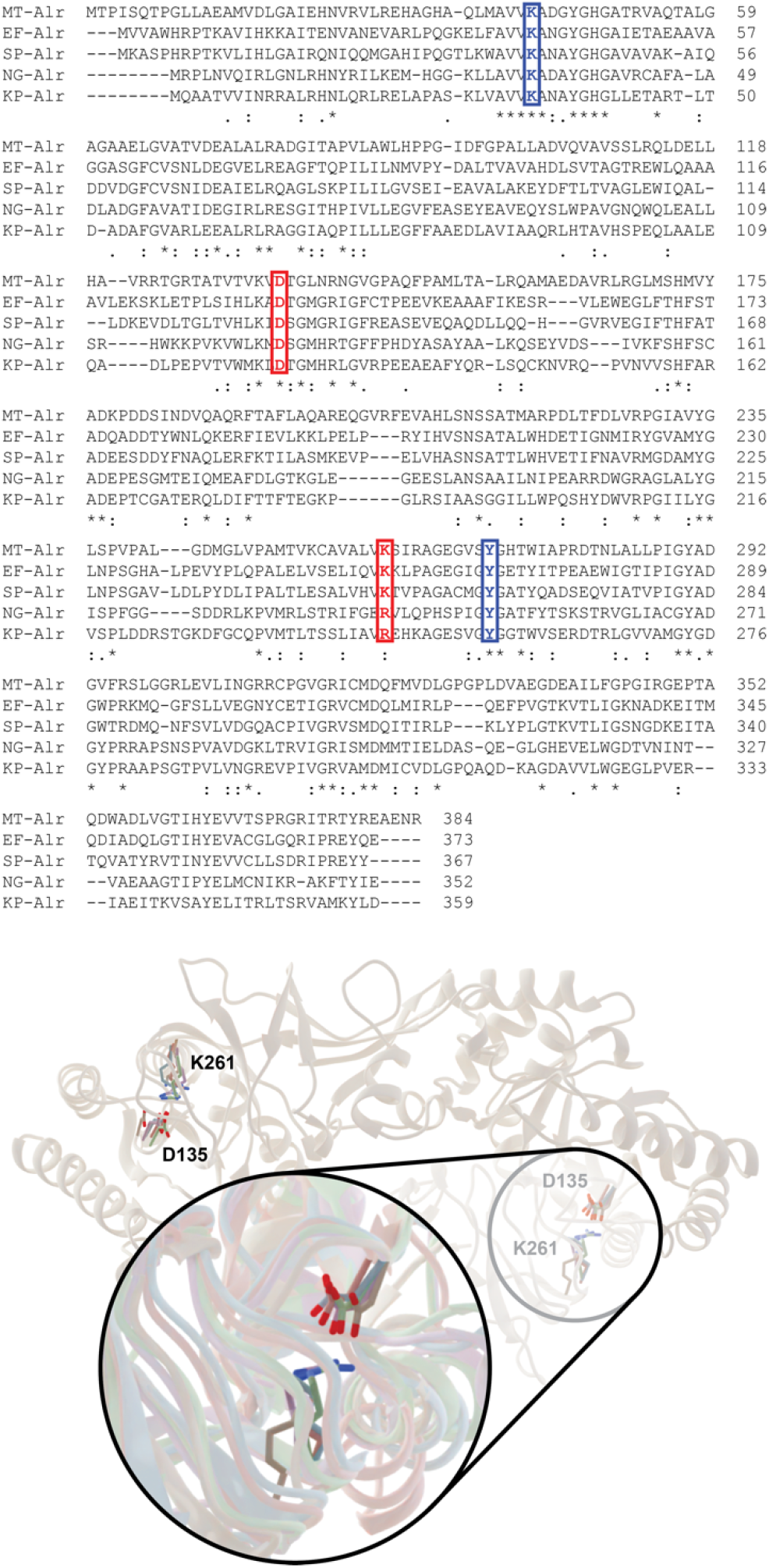
Comparison of MT-Alr and homologs. Top panel, structure-based alignment of *M. tuberculosis* (MT-Alr), *E. faecium* (EF-Alr), *S. pneumoniae* (SP-Alr), *N. gonorrhoeae* (NG-Alr), and *K. pneumoniae* (KP-Alr). Bottom panel, multiple sequence alignment of the same Alr proteins. The active site residues are boxed in blue, and the hotspot residues are in red.

## Conclusion

We report experimental and computational analyses of the Alr enzyme that suggest a novel avenue for antibacterial drug discovery. The steady-state kinetic parameters we measured for wild-type and mutant MT-Alr show that the two Asp135/Lys261 electrostatic interactions that bridge the Alr dimeric interface are critical for enzymatic activity, likely because they are required for Alr dimerization. We identified two identical pockets near each salt bridge; compounds that bind in these pockets may similarly disrupt this critical interaction. The work is significant because existing drugs bind to the orthosteric Alr active site and have unacceptable off-target effects. Our study suggests an allosteric approach to Alr inhibition that may bypass these shortcomings.

Sequence and structural alignments of several Alrs from diverse pathogenic bacteria show that this critical salt bridge is generally conserved. If the hypothesized allosteric mechanism is similarly conserved, compounds that bind in the identified pocket could enable extended-spectrum antibacterial drug discovery.

## Abbreviations

Alr: Alanine racemase
MT: *Mycobacterium tuberculosis*
PLP: Pyridoxal 5-phosphate
PCR: Polymerase chain reaction
SDS-PAGE: Sodium dodecyl sulfate-polyacrylamide gel electrophoresis
IMAC: Immobilized metal affinity chromatography
KLD: Kinase ligase DpnI enzymes
OD: Optical density

## Acknowledgements

The authors greatly acknowledge Dr. Jaeju Ko for her guidance during the initial stages of project development. The authors also acknowledge financial support from PASSHE FPDC (to SM), an IUP Senate Research Grant (to SM), and the National Institute of General Medical Sciences of the National Institutes of Health (R01GM132353 to JDD). The content is solely the responsibility of the authors and does not necessarily represent the official views of the National Institutes of Health.

## Author Contributions

**Arie Van Wieren:** Data curation; formal analysis; investigation; methodology; validation; visualization; writing review and editing. **Jacob D Durant:** Conceptualization; data curation; formal analysis; investigation; methodology; software; validation; visualization; writing original draft; writing review and editing. **Sudipta Majumdar:** Conceptualization; data curation; formal analysis; funding acquisition; investigation; methodology; supervision; project administration, software; validation; visualization; writing original draft; writing review and editing.

## References

1 Bozdogan, B. & Appelbaum, P. C. Oxazolidinones: activity, mode of action, and mechanism of resistance. Int J Antimicrob Agents 23, 113-119, 1 doi:10.1016/j.ijantimicag.2003.11.003 (2004).

2 Kim, W. et al. A selective membrane-targeting repurposed antibiotic with activity against persistent methicillin-resistant Staphylococcus aureus. Proceedings of the National Academy of Sciences 116, 16529–16534, doi:10.1073/pnas.1904700116 (2019).

3 Savoia, D. New Antimicrobial Approaches: Reuse of Old Drugs. Curr Drug Targets 17, 731–738, doi:10.2174/1389450116666150806124110 (2016).

4 Wright, G. D. Antibiotic Adjuvants: Rescuing Antibiotics from Resistance. Trends Microbiol 24, 862–871, doi:10.1016/j.tim.2016.06.009 (2016).

5 Esaki, N. & Walsh, C. T. Biosynthetic alanine racemase of Salmonella typhimurium: purification and characterization of the enzyme encoded by the alr gene. Biochemistry 25, 3261–3267, doi:10.1021/bi00359a027 (1986).

6 Walsh, C. T. Enzymes in the D-alanine branch of bacterial cell wall peptidoglycan assembly. J Biol Chem 264, 2393–2396 (1989).

7 Wang, E. & Walsh, C. Suicide substrates for the alanine racemase of Escherichia coli B. Biochemistry 17, 1313–1321, doi:10.1021/bi00600a028 (1978).

8 Nikolaidis, I., Favini-Stabile, S. & Dessen, A. Resistance to antibiotics targeted to the bacterial cell wall. Protein Science 23, 243–259, doi:10.1002/pro.2414 (2014).

9 Lambert, M. P. & Neuhaus, F. C. Mechanism of D-cycloserine action: alanine racemase from Escherichia coli W. Journal of bacteriology 110, 978–987, doi:10.1128/JB.110.3.978-987.1972 (1972).

10 Holdiness, M. R. Neurological Manifestations and Toxicities of the Antituberculosis Drugs. Medical Toxicology and Adverse Drug Experience 2, 33–51, doi:10.1007/BF03259859 (1987).

11 Yadav, S. & Rawal, G. Adverse drug reactions due to cycloserine on the central nervous system in the multidrug-resistant tuberculosis cases: a case series. PAMJ Clinical Medicine 1, 25, doi:10.11604/pamj-cm.2019.1.25.20904 (2019).

12 Schade, S. & Paulus, W. D-Cycloserine in Neuropsychiatric Diseases: A Systematic Review. The international journal of neuropsychopharmacology 19, pyv102, doi:10.1093/ijnp/pyv102 (2016).

13 Crane, G. E. The psychotropic effects of cycloserine: A new use for an antibiotic. Comprehensive Psychiatry 2, 51–59, doi:https://doi.org/10.1016/S0010-440X(61)80007-2 (1961).

14 Strych, U. & Benedik, M. J. Mutant analysis shows that alanine racemases from Pseudomonas aeruginosa and Escherichia coli are dimeric. J. Bacteriol. 184, 4321–4325, doi:10.1128/JB.184.15.4321-4325.2002 (2002).

15 Shaw, J. P., Petsko, G. A. & Ringe, D. Determination of the structure of alanine racemase from Bacillus stearothermophilus at 1.9-A resolution. Biochemistry 36, 1329–1342, doi:10.1021/bi961856c (1997).

16 LeMagueres, P. et al. Crystal structure at 1.45 A resolution of alanine racemase from a pathogenic bacterium, Pseudomonas aeruginosa, contains both internal and external aldimine forms. Biochemistry 42, 14752–14761, doi:10.1021/bi030165v (2003).

17 Ju, J. et al. Correlation between catalytic activity and monomer-dimer equilibrium of bacterial alanine racemases. Journal of biochemistry 149, 83–89, doi:10.1093/jb/mvq120 (2011).

18 Im, H., Sharpe, M. L., Strych, U., Davlieva, M. & Krause, K. L. The crystal structure of alanine racemase from Streptococcus pneumoniae, a target for structure-based drug design. BMC microbiology 11, 116, doi:10.1186/1471-2180-11-116 (2011).

19 Boggetto, N. & Reboud-Ravaux, M. Dimerization inhibitors of HIV-1 protease. Biological chemistry 383, 1321–1324, doi:10.1515/bc.2002.150 (2002).

20 Song, M., Rajesh, S., Hayashi, Y. & Kiso, Y. Design and synthesis of new inhibitors of HIV-1 protease dimerization with conformationally constrained templates. Bioorganic & medicinal chemistry letters 11, 2465–2468 (2001).

21 Strosberg, A. D. Breaking the spell: drug discovery based on modulating protein-protein interactions. Expert review of proteomics 1, 141–143, doi:10.1586/14789450.1.2.141 (2004).

22 Scheer, J. M., Romanowski, M. J. & Wells, J. A. A common allosteric site and mechanism in caspases. Proceedings of the National Academy of Sciences of the United States of America 103, 7595–7600, doi:10.1073/pnas.0602571103 (2006).

23 Van Wieren, A., Cook, R. & Majumdar, S. Characterization of Alanine Dehydrogenase and Its Effect on Streptomyces coelicolorA3(2) Development in Liquid Culture. J Mol Microbiol Biotechnol 29, 57–65, doi:10.1159/000504709 (2019).

24 Cook, R., Barnhart, R. & Majumdar, S. Effect of pH on the kinetics of alanine racemase from Mycobacterium tuberculosis. Journal of Young Investigators 36, 1–4 (2018).

25 Humphrey, W., Dalke, A. & Schulten, K. VMD: visual molecular dynamics. J Mol Graph 14, 33-38, 27-38, doi:10.1016/0263-7855(96)00018-5 (1996).

26 de Chiara, C. et al. D-Cycloserine destruction by alanine racemase and the limit of irreversible inhibition. Nat Chem Biol 16, 686–694, doi:10.1038/s41589-020-0498-9 (2020).

27 Ashkenazy, H. et al. ConSurf 2016: an improved methodology to estimate and visualize evolutionary conservation in macromolecules. Nucleic Acids Res 44, W344–350, doi:10.1093/nar/gkw408 (2016).

28 Kortemme, T., Kim, D. E. & Baker, D. Computational alanine scanning of protein-protein interfaces. Sci STKE 2004, pl2, doi:10.1126/stke.2192004pl2 (2004).

29 Durrant, J. D. & McCammon, J. A. BINANA: a novel algorithm for ligand-binding characterization. J Mol Graph Model 29, 888–893, doi:10.1016/j.jmgm.2011.01.004 (2011).

30 Meng, E. C., Pettersen, E. F., Couch, G. S., Huang, C. C. & Ferrin, T. E. Tools for integrated sequence-structure analysis with UCSF Chimera. BMC Bioinformatics 7, 339, doi:10.1186/1471-2105-7-339 (2006).

31 LeMagueres, P. et al. The 1.9 A crystal structure of alanine racemase from Mycobacterium tuberculosis contains a conserved entryway into the active site. Biochemistry 44, 1471–1481, doi:10.1021/bi0486583 (2005).

32 Jacobson, M. P. et al. A hierarchical approach to all-atom protein loop prediction. Proteins 55, 351–367, doi:10.1002/prot.10613 (2004).

33 Jacobson, M. P., Friesner, R. A., Xiang, Z. & Honig, B. On the role of the crystal environment in determining protein side-chain conformations. J Mol Biol 320, 597–608, doi:10.1016/s0022-2836(02)00470-9 (2002).

34 Sastry, G. M., Adzhigirey, M., Day, T., Annabhimoju, R. & Sherman, W. Protein and ligand preparation: parameters, protocols, and influence on virtual screening enrichments. J Comput Aided Mol Des 27, 221–234, doi:10.1007/s10822-013-9644-8 (2013).

35 Halgren, T. A. Identifying and characterizing binding sites and assessing druggability. J Chem Inf Model 49, 377–389, doi:10.1021/ci800324m (2009).

36 Kozakov, D. et al. The FTMap family of web servers for determining and characterizing ligand-binding hot spots of proteins. Nat Protoc 10, 733–755, doi:10.1038/nprot.2015.043 (2015).

37 Kochnev, Y. & Durrant, J. D. FPocketWeb: protein pocket hunting in a web browser. J Cheminform 14, 58, doi:10.1186/s13321-022-00637-0 (2022).

38 Le Guilloux, V., Schmidtke, P. & Tuffery, P. Fpocket: an open source platform for ligand pocket detection. BMC Bioinformatics 10, 168, doi:10.1186/1471-2105-10-168 (2009).

39 Schmidtke, P., Le Guilloux, V., Maupetit, J. & Tuffery, P. fpocket: online tools for protein ensemble pocket detection and tracking. Nucleic Acids Res 38, W582–589, doi:10.1093/nar/gkq383 (2010).

40 Cardinale, D. et al. Homodimeric enzymes as drug targets. Current medicinal chemistry 17, 826–846 (2010).

41 Awasthy, D., Bharath, S., Subbulakshmi, V. & Sharma, U. Alanine racemase mutants of Mycobacterium tuberculosis require D-alanine for growth and are defective for survival in macrophages and mice. Microbiology (Reading) 158, 319–327, doi:10.1099/mic.0.054064-0 (2012).

42 Ciustea, M. et al. Thiadiazolidinones: a new class of alanine racemase inhibitors with antimicrobial activity against methicillin-resistant Staphylococcus aureus. Biochem Pharmacol 83, 368–377, doi:10.1016/j.bcp.2011.11.021 (2012).

43 Jyothikumar, J., Chandani, S. & Ramakrishna, T. Computational evidence of a new allosteric communication pathway between active sites and putative regulatory sites in the alanine racemase of Mycobacterium tuberculosis. bioRxiv, 346130 (2018).

