## supporting information for "Computational and experimental analyses of alanine racemase suggest new avenues for developing allosteric small-molecule antibiotics"

[3] Current address: The Johns Hopkins University School of Medicine, Baltimore, MD 21205

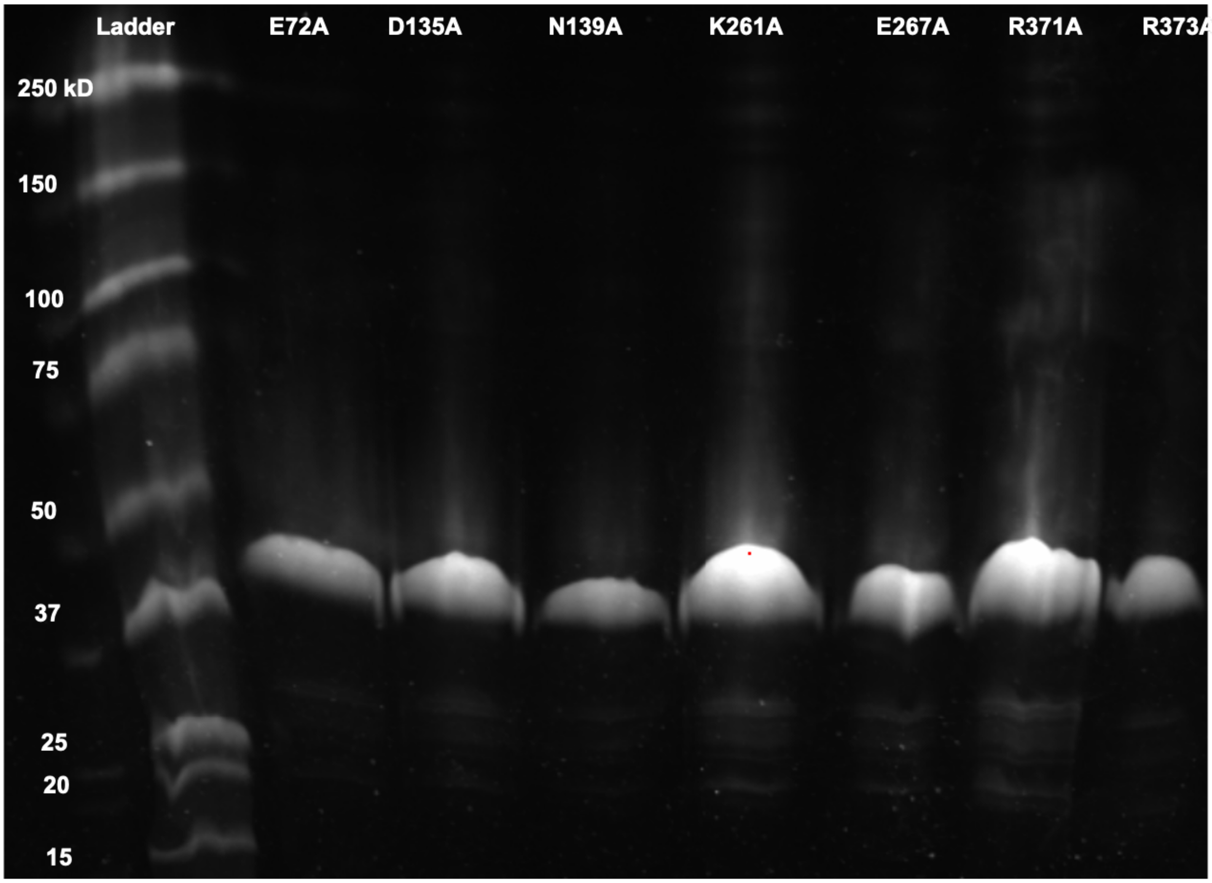

Fig S1. SDS-PAGE of purified MT-Alr mutant proteins.

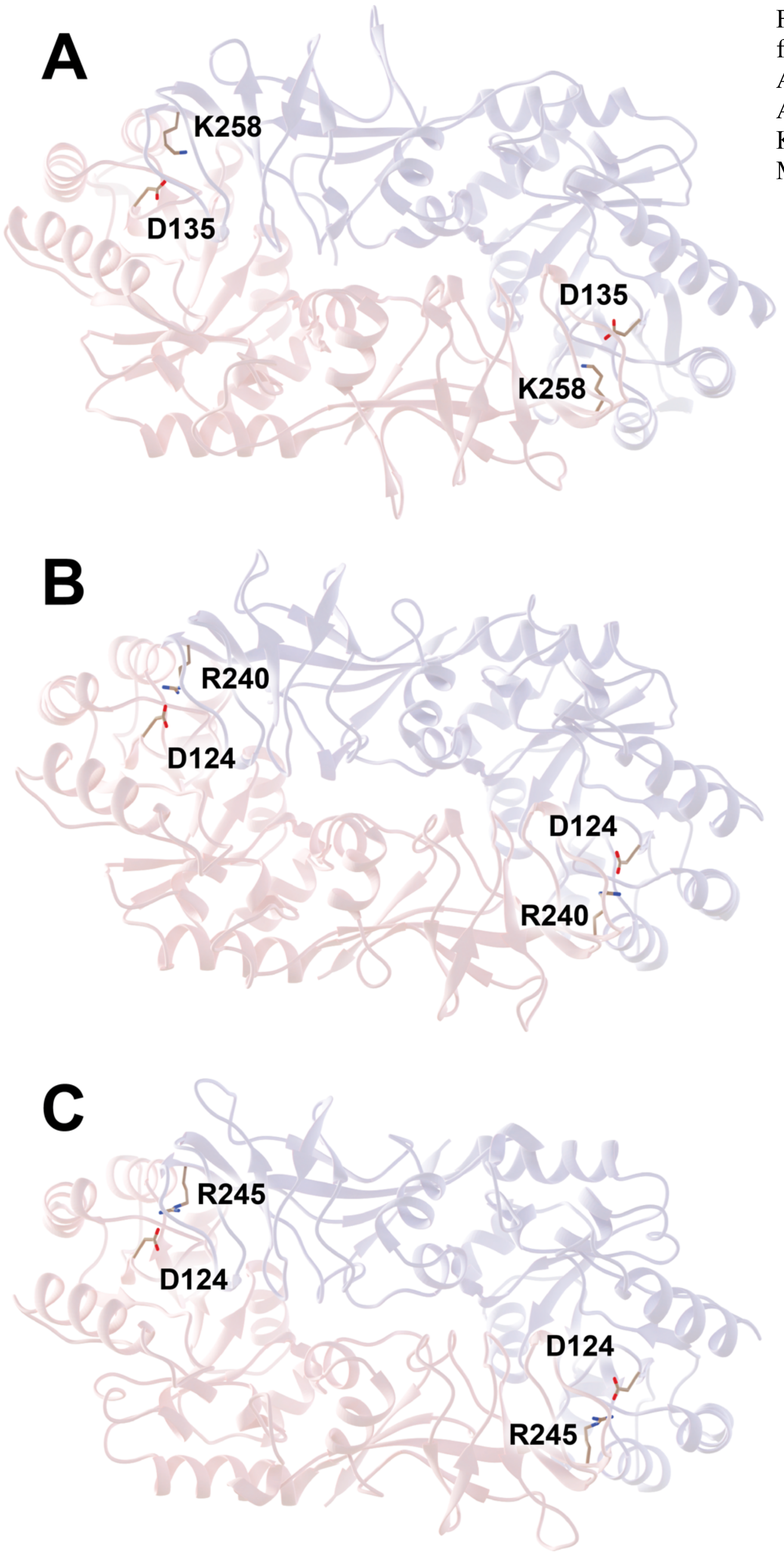


Fig S2. Homology models of Alrs from (A) *Enterococcus faecium*, EF-Alr; (B) *Neisseria gonorrhoeae*, NG-Alr; and (C) *Klebsiella pneumoniae*, KP-Alr, generated using SWISS-MODEL.

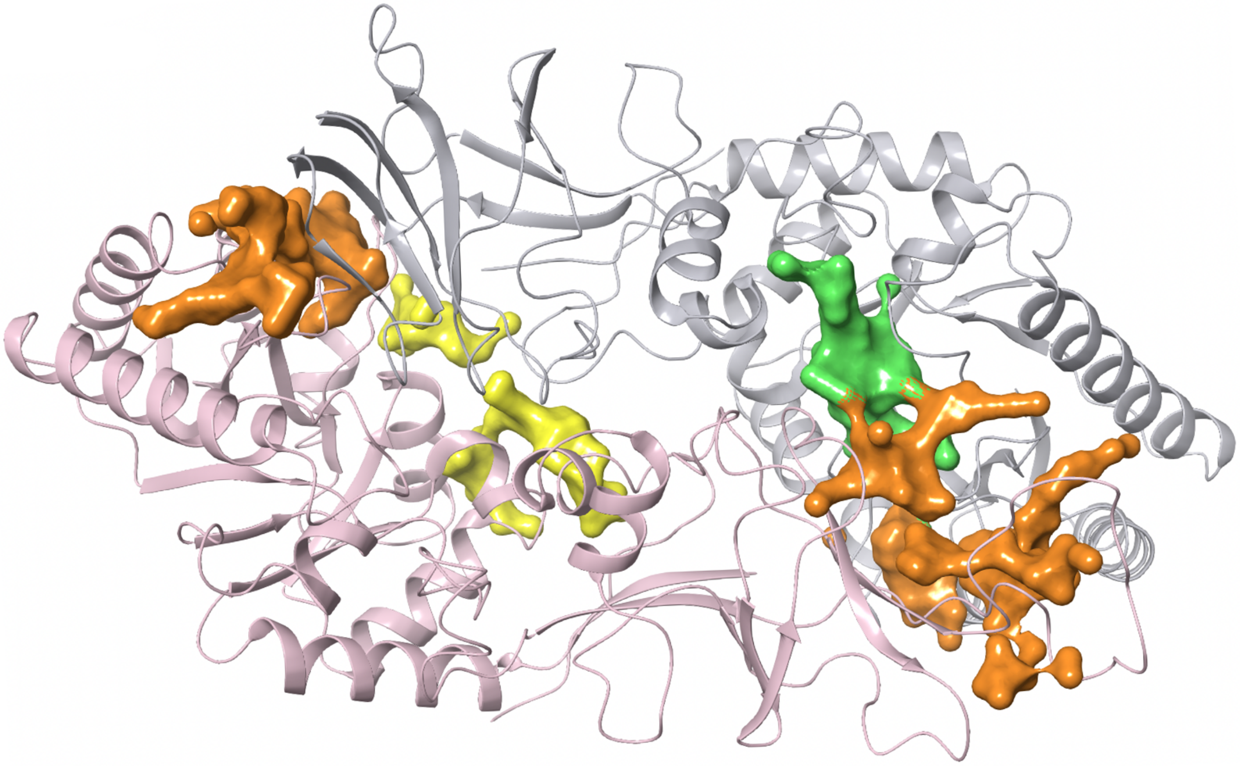

Fig S3. SiteMap-identified pockets near the Asp135/Lys261 salt bridge (orange), orthosteric cycloserine-binding site (green), and central Arg371/Arg371 juxtaposition (yellow).

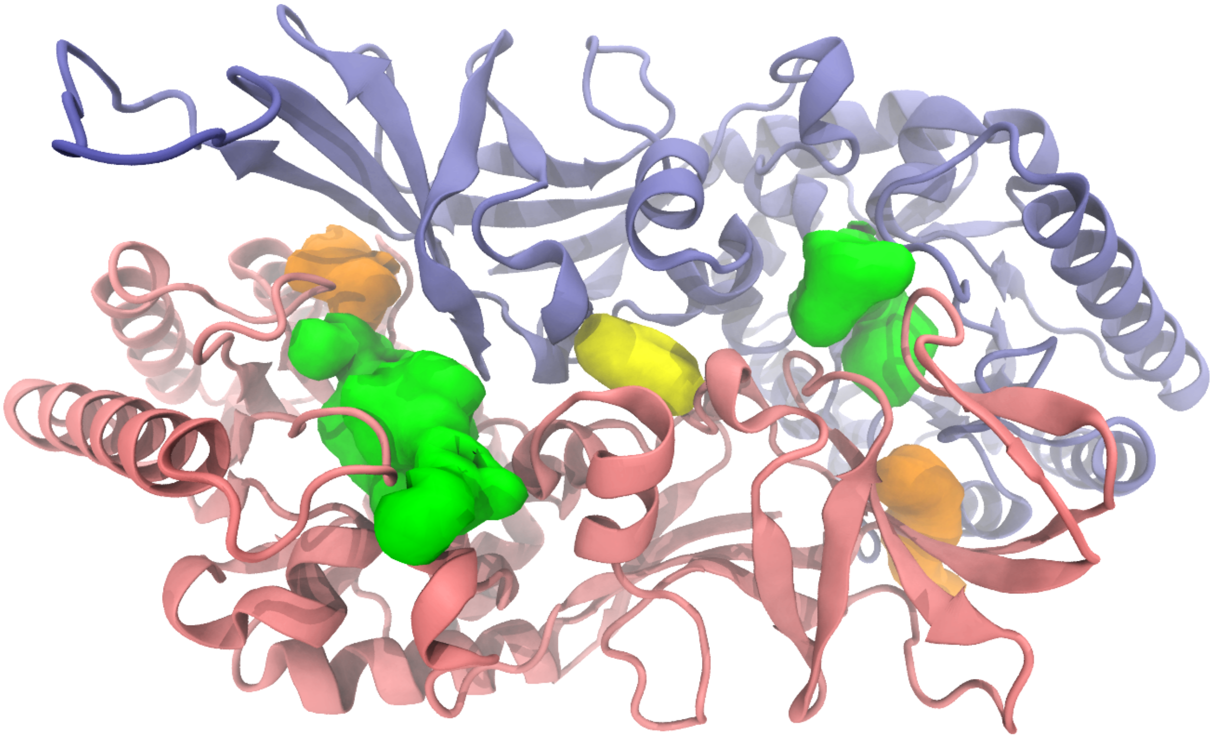

Fig S4. FPocketWeb-identified pockets near the Asp135/Lys261 salt bridge (orange), orthosteric cycloserine-binding site (green), and central Arg371/Arg371 juxtaposition (yellow).

Scheme S1. The MT-Alr-catalyzed isomerization of D-alanine to L-alanine enables the subsequent $NAD^+$-dependent oxidation of L-alanine to pyruvate by alanine dehydrogenase (Ald).

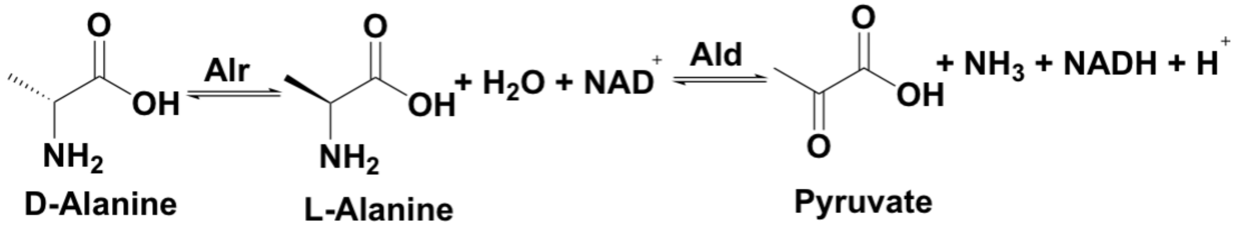
